# The experimental determination of redox-potential of mutated cytochrome *c* with eight charge changing substitutions

**DOI:** 10.1101/015131

**Authors:** V.V. Ptushenko, R.V. Chertkova

## Abstract

The redox potential of the mutated cytochrome *c* form with eight charge changing substitutions was experimentally determined.

## Introduction

Cytochrome *c* (cyt *c*) is one of the key proteins in cell respiration, as well as in some pathological processes such as ROS generation or apoptosis development (Skulachev, 1998). We have previously investigated a number of mutated forms of horse heart cyt *c*, including the variant with eight charge changing substitutions (mut8), K8E/K27E/E62K/E69K/K72E/K86E/K87E/E90K, and it was shown that it demonstrated significantly lower rates of reactions with components of respiration chain, ubiquinol:cytochrome *c* reductase (complex III) and cytochrome *c* oxidase (complex IV) than wild-type cyt *c* (Pepelina et al., 2009). This decrease in reaction rates can in principle be related both to a violation of docking and to the change in the redox potential *E*_m_ of cyt *c* heme, which would result from charge changing substitutions in its amino acid composition. The aim of this work was to determine experimentally the *E*_m_ of the mutated form mut8 of cyt *c*.

## Methods

The redox titration of cyt *c* was performed under anaerobic conditions and with continuous stirring of solution in an optical cell of Varian “Care 50” spectrophotometer (USA). The ratio of the oxidized and reduced forms of cyt *c* was determined by absorption spectrum, according to *A*_549_ – *A*_556_. The medium contained 1 mM cyt *c*, 20 mM Tris-HCl (pH 7.5), 10 mM diaminodurol, 100 mM KCl. Oxidation or reduction of cyt *c* was carried out by adding a solution of ferricyanide or ascorbate, respectively. The redox potential of the solution was measured using a Pt, Ag/AgCl electrode pair.

## Results and Discussion

Fig. 1 shows the redox titration curves of cyt *c*, WT and mut8. It can be seen that the redox potential of the mutated cyt *c* form is 27 meV higher than that of WT cyt *c*. This mutated form differs from the wild type form by 8 amino acid residues substitutions, each of them is entered instead of a charged WT residue and carries an opposite charge. Thus, the five lysine residues (K8, K27, K72, K86 and K87) are substituted for glutamate residues. At the same time, the three glutamate residues (E62, E69 and E90) are replaced by lysine residues. Since the replacement of positive charged lysine residues to negative charged glutamate residues and inverse substitution do not completely compensate each other, the cyt *c* must acquire some excessive negative charge, which would lead to some negative shift in cyt *c E*_m_. Our experimental data confirms this expectation.

**Figure 1.**
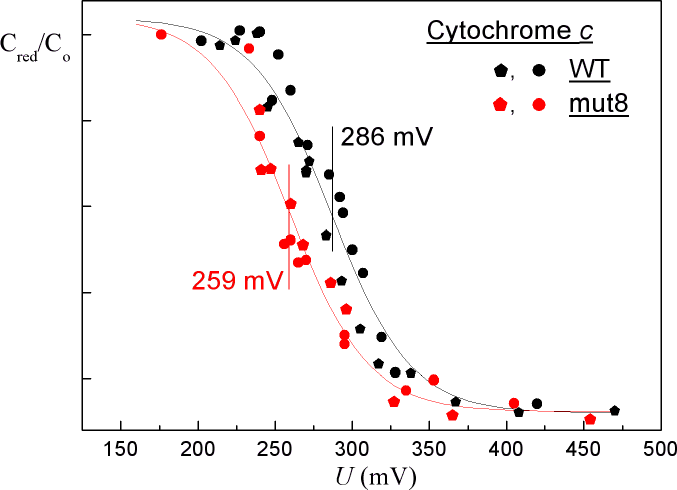
The redox titration curves of cyt *c*, wild type (WT; black symbols) and mutated (mut8; red symbols) forms. Circles and polygons correspond to different replicates of titration.

## Acknowledgements

We express our thanks to Dr. A.Y. Semenov whose insistence and chivalry made this work possible.

